# Transient T cell expansion, activation, and proliferation in therapeutically vaccinated SIV+ macaques treated with N-803

**DOI:** 10.1101/2022.07.08.499199

**Authors:** Olivia E. Harwood, Alexis J. Balgeman, Abigail J. Weaver, Amy L. Ellis-Connell, Andrea M. Weiler, Katrina N. Erickson, Lea M. Matschke, Athena E. Golfinos, Vaiva Vezys, Pamela J. Skinner, Jeffrey T. Safrit, Paul T. Edlefsen, Matthew R. Reynolds, Thomas C. Friedrich, Shelby L. O’Connor

## Abstract

Vaccine strategies aimed at eliciting HIV-specific CD8+ T cells are one major target of interest in HIV functional cure strategies. We hypothesized that CD8+ T cells elicited by therapeutic vaccination during antiretroviral therapy (ART) would be recalled and boosted by treatment with the IL-15 superagonist N-803 after ART discontinuation. We intravenously immunized four SIV+ Mauritian cynomolgus macaques (MCM) receiving ART with vesicular stomatitis virus (VSV), modified vaccinia virus Ankara strain (MVA), and recombinant adenovirus serotype 5 (rAd-5) vectors all expressing SIVmac239 Gag. Immediately after ART cessation, these animals received three doses of N-803. Four control animals received no vaccines or N-803. The vaccine regimen generated a high magnitude of Gag-specific CD8+ T cells that were proliferative and biased toward an effector memory phenotype. We then compared cells elicited by vaccination (Gag-specific) to cells elicited by SIV infection and unaffected by vaccination (Nef-specific). We found that N-803 treatment enhanced both the frequencies of bulk and proliferating antigen-specific CD8+ T cells elicited by vaccination and the antigen-specific CD8+ T cells elicited by SIV infection. In sum, we demonstrate that a therapeutic heterologous prime-boost-boost (HPBB) vaccine can elicit antigen-specific effector memory CD8+ T cells that are boosted by N-803.

## Introduction

CD8+ T cells play a critical role in controlling HIV/SIV replication through cytolytic and non-cytolytic means, both in the presence or absence of ART (1, 2). Indeed, spontaneous control of HIV/SIV replication without ART, known as elite control (EC), is associated with the higher frequency and prolonged survival of CD8+ T cells, and non-cytolytic virus inhibition by CD8+ T cells (3, 4). While HIV and SIV replication can be durably suppressed by antiretroviral therapy (ART), under most circumstances virus replication rebounds when ART is discontinued (5, 6). Thus, a core strategy of HIV cure initiatives is inducing potent CD8+ T cells through therapeutic vaccinations to enable people with HIV to stop ART and durably control virus replication.

Expansion of CD8+ T cells during acute HIV/SIV infection is associated with peak viral load decline (7–9). During the progression of untreated infection, however, chronic antigen exposure leads to the loss of CD8+ T cell function and increased CD8+ T cell apoptosis (4, 10). Potent CD8+ T cell responses can also put pressure on viral evolution and, unfortunately, lead to viral escape (11, 12). One major challenge for therapeutic vaccines is to supersede naturally elicited cellular immunity, suppress virus replication without ART, and avoid the selection of viral variants.

Many current HIV vaccine strategies can induce neutralizing antibodies and/or CD8+ T cells, but it is essential to further understand the features of protective cellular immune responses and determine how best to elicit those responses (13–17). Heterologous prime-boost (HPB) vaccines induce superior cellular immunity compared to single-dose strategies and homologous prime-boost strategies (18, 19). HPB vaccine strategies elicit abundant CD8+ T cells with enhanced virus suppression, proliferative capabilities, and long-term survival. These vaccines also expand the populations of central memory (T_CM_) and effector memory (T_EM_) T cells (15, 19–22). Antigen-specific CD8+ T cells with T_EM_ phenotypes have exhibited high proliferative capacities, inhibitory effects against HIV, and have specifically been implicated in EC (13, 21).

Numerous therapeutic vaccine regimens have demonstrated the potential to elicit anti-SIV cellular immunity in macaque models, particularly both T_CM_ and T_EM_ responses (23–25). Transient antigen exposure provided by nonreplicating vectors favors the induction of T_CM_ responses. T_CM_ expansion and production of effector cells are important for the anti-HIV immune response, but T_CM_ populations require time to respond via anamnestic expansion upon antigen recognition (26). Replicating vectors that provide recurrent antigen exposure, on the other hand, favor the induction of protective T_EM_ responses, which can deliver prompt effector function without antigen re-stimulation (27).

Therapeutic HIV vaccines could improve the host antiviral immune response and benefit those already infected with HIV (28). In ART treated SIV+ macaques, several vaccine modalities have induced high-frequency cellular immune responses against multiple SIV proteins (i.e. Gag, Pol, and Env) (23, 29, 30). High magnitude Gag-specific responses have also been detected in therapeutic vaccine strategies delivering Gag immunogens only (24, 25). T cell recognition of multiple epitopes in Gag has been associated with lower viremia, suggesting Gag may be a promising immune target for vaccine strategies (24, 31–33). Some vaccine strategies have conferred improved viral control after suspending ART, but success is limited (23, 24, 29). This may be a consequence of eliciting too few functional antigen-specific T cells in the correct locations to suppress the rebounding virus when ART is interrupted. A primary objective of therapeutic vaccines is to educate the host cellular immune system to develop characteristics associated with EC, such as T_EM_ phenotype, proliferative capacity, high magnitude, and antigen specificity. A heterologous prime-boost-boost (HPBB) strategy is one novel approach that may achieve this goal.

Cytokine administration can be used as an adjuvant for T cell-based therapies (34, 35). The proinflammatory cytokine IL-15 is a critical regulator and promoter of memory T cell development and maintenance (36, 37). IL-15 is involved in NK cell and T cell generation, homeostasis, maturation, and support (38, 39). N-803 (formerly Alt-803) is an IL-15 superagonist complex exhibiting superior half-life, bioactivity, efficacy, and tissue retention compared to IL-15. N-803 has been shown to increase the frequency, activation, proliferation, and function of CD4+ T cells, CD8+ T cells, and NK cells in mice and macaques (40–42). In SIV+ macaques, N-803 can activate and expand CD8+ T cells and NK cells and direct these cells to the lymph nodes (LNs) (43–45). However, the impact of N-803 on CD8+ T cells elicited by therapeutic HIV/SIV vaccines is unknown.

In the present study, we hypothesized that combining HPBB vaccination with N-803 treatment could elicit and recall a high magnitude of Gag-specific CD8+ T cells and target them to the LNs. We vaccinated four SIV+ ART-treated Mauritian cynomolgus macaques (MCM) with an HPBB regimen consisting of a vesicular stomatitis virus (VSV) prime followed by two boosts with modified vaccinia virus Ankara strain (MVA) and recombinant adenovirus serotype 5 (rAd-5), all expressing SIVmac239 Gag. Four unvaccinated SIV+ ART-treated macaques served as controls. Six weeks after the third vaccine, we discontinued ART for all eight animals. The vaccinated animals received three doses of N-803. We report that this therapeutic regimen enhanced the frequency of Gag-specific lymphocytes with phenotypes associated with activation (CD69), proliferation (Ki-67), and memory (T_EM_) in the peripheral blood and LNs of the vaccinated animals.

## Results

### Animal study design, infection, animals, and viral loads

We infected eight Mauritian cynomolgus macaques (MCMs) i.v. with SIVmac239M (Figure 1A). All animals expressed at least one copy of the M3 MHC haplotype that is associated with a high SIV viral load set point, and none possessed the M1 haplotype associated with SIV control (Table 1) (46, 47). Animals exhibited peak viremia between 3×10^5^ and 4×10^7^ ceq/mL on day 11 post-infection (Figure 1B). Animals began a daily ART regimen of dolutegravir (DTG), tenofovir disoproxil fumarate (TDF), and emtricitabine (FTC) at 14 days post-infection. Fourteen days was chosen because, in rhesus macaque studies, this allows time for seroconversion, reservoir establishment, and the generation of a T cell response prior to virus suppression, but not so much time to enable viral immune escape and immune exhaustion (48, 49). Viremia was suppressed below the limit of detection in all 8 animals within 42 days of starting ART. Four animals (cy1035, cy1036, cy1039, and cy1043) were therapeutically vaccinated i.v. with an HPBB vaccine strategy under the control of ART, beginning ∼4.5 months after infection (Figure 1A, blue animals). Animals first received VSV expressing SIVmac239 Gag, followed by boosts with MVA expressing SIVmac239 Gag-Pol and rAd-5 expressing SIVmac239 Gag. Inoculations were separated by 4 weeks. These vectors have been shown to elicit robust antigen-specific CD8+ T cells (15, 29). Four animals (cy1037, cy1040, cy1044, and cy1045) were left unvaccinated as controls and received saline injections, rather than vaccine vectors (Figure 1A, red animals).

**Figure 1.**
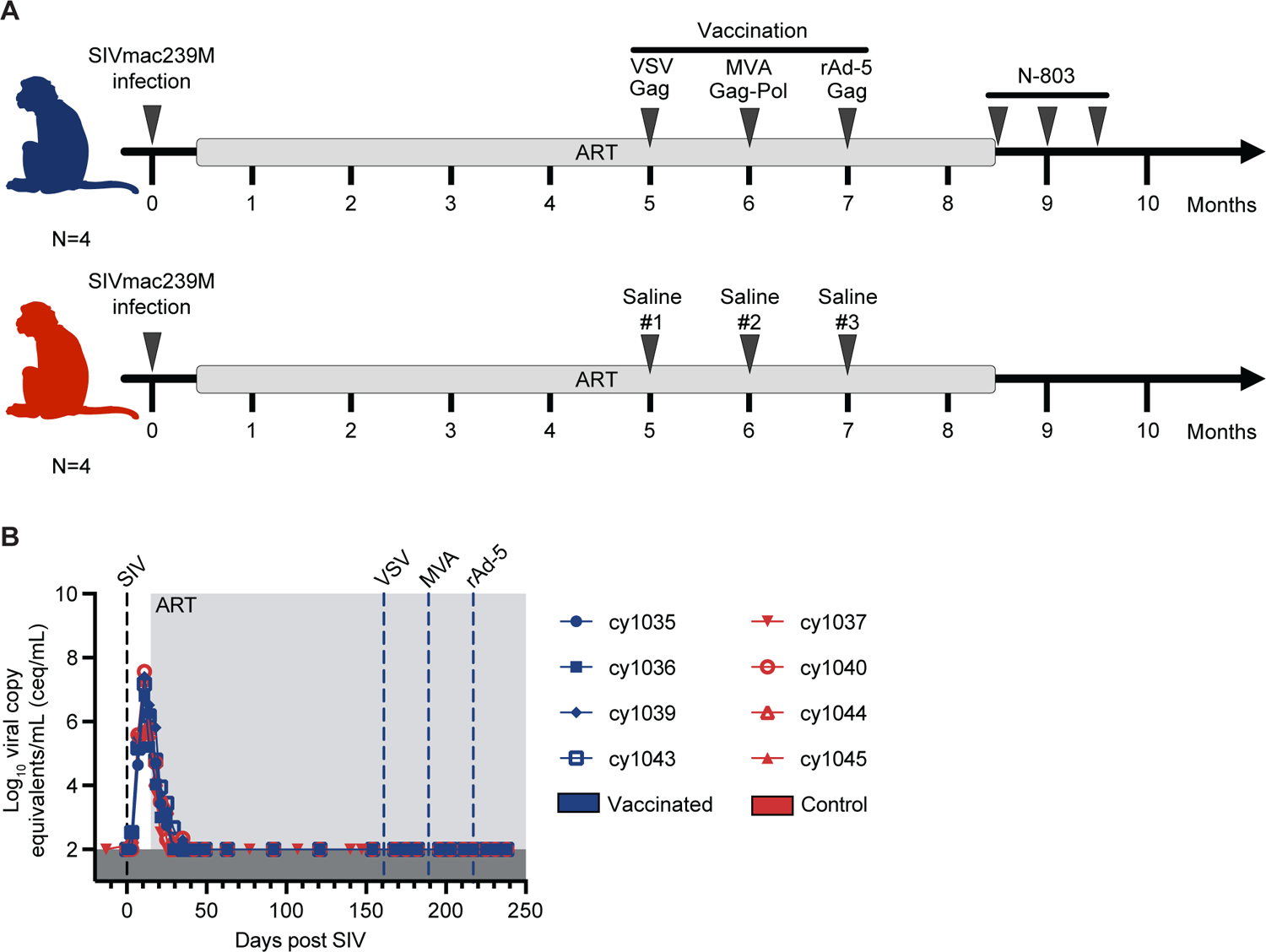
Experimental design and longitudinal plasma viral loads. (A) Study design depicting timeline of SIV infection, therapeutic vaccination, ART release, and N-803 delivery: eight male MCM were infected i.v. with 10,000 IU of SIVmac239M. All eight began receiving ART 14 dpi. Four MCM (blue, vaccinated animals) were sequentially immunized with recombinant heterologous viral vectors VSV, MVA, and rAd-5 each encoding SIVmac239 Gag. ART was discontinued in all eight animals six weeks after the final boost. The vaccinated animals, but not the control animals, received 3 doses of N-803 beginning three days after ART release. **(B)** Individual plasma viral loads from SIV inoculation through vaccination. Viral loads are displayed as log_10_ceq/mL.

**TABLE 1.**
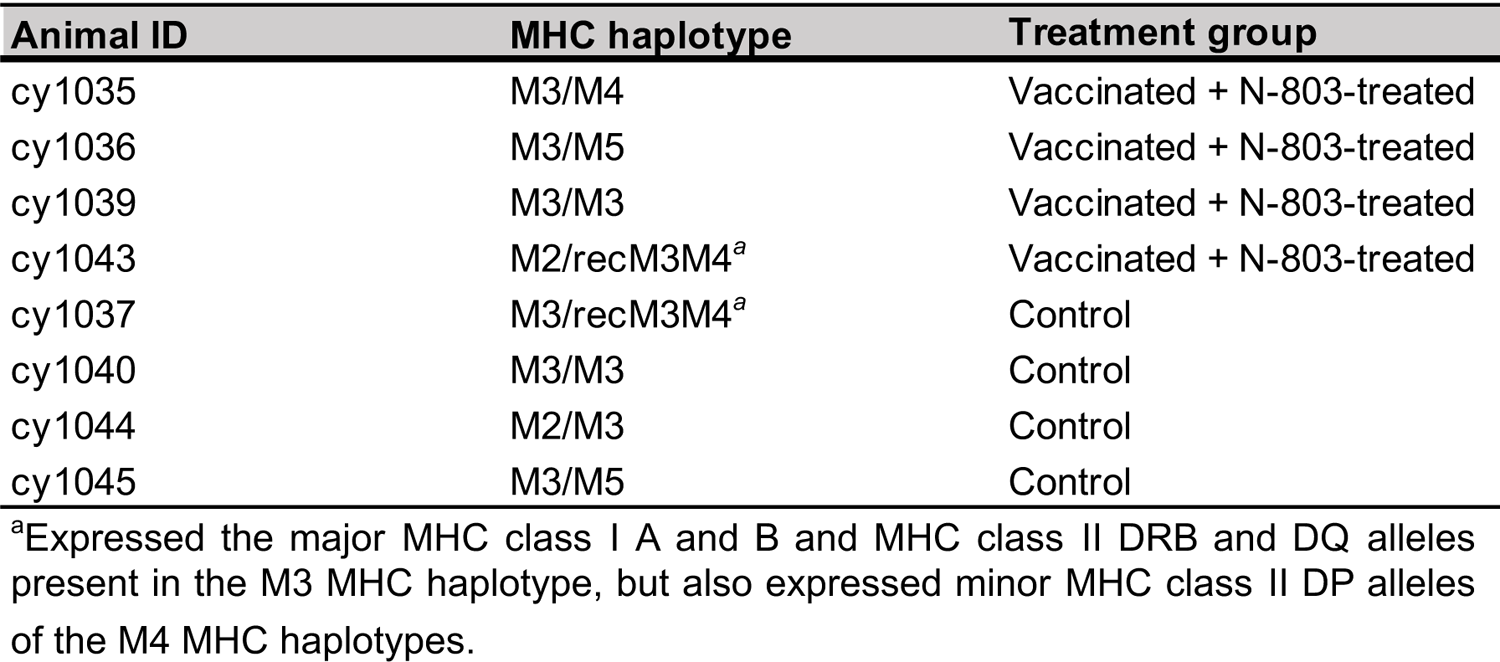
Animals used in this study

| Animal ID | MHC haplotype | Treatment group |
| --- | --- | --- |
| cy1035 | M3/M4 | Vaccinated + N-803-treated |
| cy1036 | M3/M5 | Vaccinated + N-803-treated |
| cy1039 | M3/M3 | Vaccinated + N-803-treated |
| cy1043 | M2/recM3M4<sup>a</sup> | Vaccinated + N-803-treated |
| cy1037 | M3/recM3M4<sup>a</sup> | Control |
| cy1040 | M3/M3 | Control |
| cy1044 | M2/M3 | Control |
| cy1045 | M3/M5 | Control |

<sup>a</sup>Expressed the major MHC class I A and B and MHC class II DRB and DQ alleles present in the M3 MHC haplotype, but also expressed minor MHC class II DP alleles of the M4 MHC haplotypes.

### Therapeutic i.v. HPBB vaccination elicits robust and broad Gag-specific T cell responses

To assess the immunogenicity of the HPBB vaccination, we performed IFN-γ ELISPOT assays using PBMCs collected before the first inoculation and 7 days after each vaccination, as well as 21 days after the rAd-5 vaccination. We stimulated the PBMCs with Mafa-A1*063-restricted Gag_386-394_GW9 or Nef_103-111_RM9 peptides, or a Gag peptide pool spanning the SIVmac239 Gag proteome. PBMCs from vaccinated animals responded to stimulation with the Gag pool and Gag_386-394_GW9 peptide throughout the vaccine phase (Figure 2A, left). The immunodominant CD8+ T cell response targeting the Gag_386-394_GW9 epitope was consistent with studies of other SIV+ *Mafa-A1*063+* MCMs (47, 49). As anticipated, the CD8+ T cell responses against the Gag pool or Gag_386-394_GW9 were substantially lower in the control group (Figure 2A, right). CD8+ T cell responses specific for the Nef_103-111_RM9 peptide were low across both groups of animals. Further, the Gag_386-394_GW9 IFN-γ ELISPOT responses in the vaccinated animals were ∼10-50-fold greater than the Nef_103-111_RM9 responses while the Gag_386-394_GW9 responses in the control animals were approximately the same as the Nef_103-111_RM9 responses (Figure 2A). These data demonstrate the difference between the frequency of vaccine-elicited versus naturally elicited antigen-specific T cells in ART-suppressed SIV+ animals.

**Figure 2.**
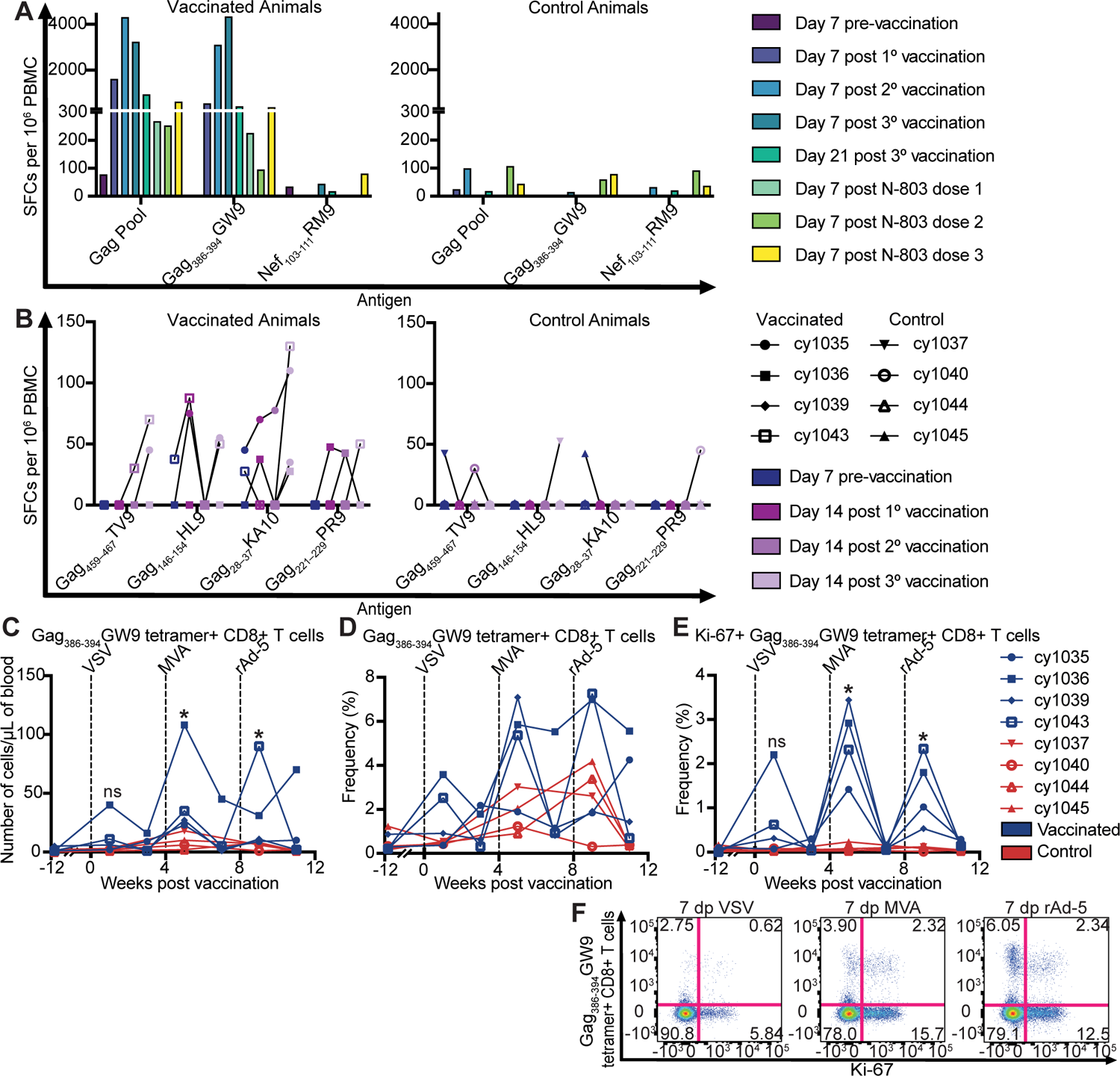
High magnitude, breadth, and proliferation of Gag–specific CD8+ T cell responses in the PBMC induced by HPBB vaccination. **(A)** IFN-γ ELISPOT assays were performed longitudinally at the indicated time points to assess responses to Gag_386-394_GW9, Nef_103-111_RM9, and a Gag peptide pool spanning the SIVmac239 Gag proteome in the vaccinated (left) and control (right) animals. Bars are displayed as the median. **(B)** IFN-γ ELISPOT assays were performed longitudinally at the indicated time points to assess responses to Gag_459-467_TV9, Gag_146-154_HL9, Gag_28-37_KA10, and Gag_221-229_PR9 in the vaccinated (left) and control (right) animals. Results are displayed for each individual animal. **(C)** Number of Gag_386-394_GW9 tetramer+ CD8+ T cells per μL of blood throughout the vaccine phase in the vaccinated (blue) and control (red) animals. **(D)** Frequency of Gag_386-394_GW9 tetramer+ CD8+ T cells throughout the vaccine phase in the vaccinated (blue) and control (red) animals. **(E)** Frequency of CD8+ T cells that are both Gag GW9-specific and Ki-67+ throughout the vaccine phase in the vaccinated (blue) and control (red) animals. **(F)** Flow cytometry dot plots illustrating expansion of proliferating (Ki-67+) Gag GW9-specific CD8+ T cells after VSV-, MVA-, and rAd-5-SIVmac239 Gag immunizations. Results are displayed for each animal individually. * *P*=0.0286. *P* values were calculated using Mann-Whitney U tests comparing the vaccinated and control groups at each indicated point.

Next, we used cryopreserved PBMCs to assess T cell responses against SIV epitopes restricted by other MHC class I alleles expressed by the M3 MHC haplotype (Gag_459-467_TV9, Gag_146-154_HL9, Gag_28-37_KA10, and Gag_221-229_PR9) (50). Some control animals exhibited low-level (≤50 SFCs per 10^6^ PBMC after background subtraction) responses to Gag_459-467_TV9, Gag_146-154_HL9, Gag_28-37_KA10, and Gag_221-229_PR9, but these were independent of time point and did not uniformly increase throughout infection (Figure 2B, right). The vaccinated animals exhibited a slightly higher frequency of IFN-γ+ responses (50-150 SFCs per 10^6^ PBMC) to Gag_459-467_TV9, Gag_146-154_HL9, Gag_28-37_KA10, and Gag_221-229_PR9 than the control animals throughout the vaccine phase, suggesting that vaccination elicited a population of T cells targeting a greater diversity of Gag peptides (Figure 2B, left).

### Therapeutic i.v. HPBB vaccination increases the frequency of Gag-specific CD8+ T cells

We next assessed the frequency and phenotypes (Supplemental Figure 1) of Gag_386-394_GW9 tetramer+ CD8+ T cells (henceforth referred to as Gag GW9-specific CD8+ T cells) longitudinally in PBMC collected prior to and during vaccination. The number of Gag GW9-specific CD8+ T cells was significantly higher (marked by an asterisk) in the vaccinated animals after the MVA and rAd-5 vaccines when compared to the control animals (Figure 2C). The frequency of Gag GW9-specific CD8+ T cells in some of the vaccinated animals was higher after each vaccine than in the control animals, but this was not statistically significant, likely because the vaccines also drastically increased the frequency of bulk CD8+ T cell populations (Figure 2D and Supplemental Figure 2). We also compared the frequency of Ki-67+ and Gag GW9 tetramer double-positive CD8+ T cells (Figure 2F, top right quadrants) between the vaccinated and control animals after each vaccine. The frequency of these cells was significantly higher in the vaccinated animals after the MVA and rAd-5 vaccines when compared to control animals (Figure 2E). The Ki-67+ Gag GW9-specific CD8+ T cells remained less than 0.25% in the control animals throughout the entire study (Figure 2E, red animals). Similarly, the frequency of bulk CD8+ T cells and CD8+ T_EM_ expressing the proliferation marker Ki-67 alone or in combination with the activation marker CD69 increased in the vaccinated animals one week after each of the three vaccines, and the frequency of bulk CD4+ T cells expressing Ki-67 or Ki-67 and CD69 increased in the vaccinated animals one week after the second vaccine (Supplemental Figure 2, blue animals).

### T_EM_ phenotype bias in Gag-specific CD8+ T cells elicited by therapeutic i.v. HPBB vaccination

The frequencies of Gag GW9-specific CD8+ T_CM_ and T_EM_ cells (Figure 3A) were compared between the vaccinated and control animals seven days after each of the three vaccinations. The frequency of both Gag GW9-specific T_CM_ and T_EM_ cells were similar between the treatment and control animals on day seven post-VSV or saline #1 (Figure 3B). However, on day seven after the second and third injections, we detected an increased frequency of Gag GW9-specific T_EM_ cells in the vaccinated animals, while the proportion of Gag GW9-specific T_EM_ cells declined in the control group. These results suggest that the vaccine regimen biased antigen-specific CD8+ T cells to an effector memory phenotype.

**Figure 3.**
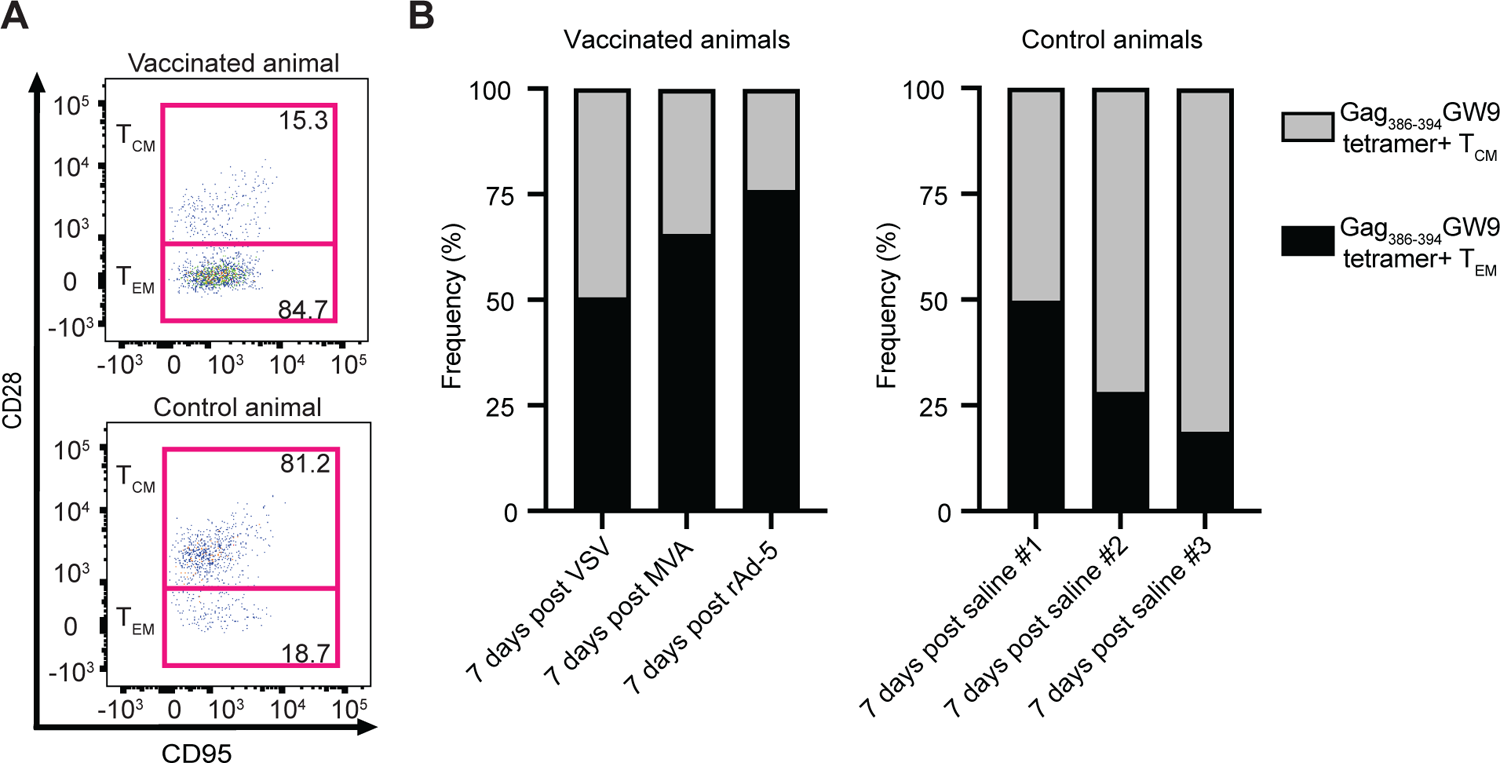
Gag GW9-specific CD8+ T_CM_ and T_EM_ cells in the PBMC throughout vaccination. **(A)** Representative flow cytometry dot plots from a vaccinated animal (top plot) and a control animal (bottom plot) illustrating the frequency of Gag GW9-specific CD8+ T_CM_ cells (top gate, CD28+CD95+) and T_EM_ cells (bottom gate, CD28-CD95+) after rAd-5-SIVmac239 Gag immunization. **(B)** Stacked bar plots showing the mean frequencies of Gag GW9-specific T_CM_ cells (gray) and Gag GW9-specific T_EM_ cells (black) in the peripheral blood of the vaccinated (left) and control (right) animals at day 7 post each vaccination.

### Gag-specific activation-induced marker (AIM) expression increases following therapeutic i.v. HPBB vaccination

To further characterize the functional profile of vaccine-elicited Gag-specific T cells, we used the AIM assay to measure the expression of effector molecules by CD4+ and CD8+ T cells after in vitro Gag stimulation. In this assay, we used cells collected before the first vaccination and three weeks after the last vaccination (Figure 4) and quantified the expression of CD25 and CD69 that indicate early activation, CD137 that marks antigen recognition, and CD107a that is associated with cytokine secretion and degranulation (51–55). Prior to vaccination, there were no significant differences in the frequency of cells expressing activation markers between unstimulated and Gag-stimulated conditions (Figure 4, left). The frequency of cells producing activation markers in the unstimulated conditions was generally higher before vaccination when compared to after vaccination. We suspect this is due to residual immune activation from acute SIV infection (56). Post-vaccination, however, we observed distinct populations of CD4+ and CD8+ T cells expressing activation markers after being stimulated with the Gag peptide pool (Figure 4, right). These results support that the vaccine regimen expanded the repertoire of Gag-specific T cells with the potential to respond to antigen and exhibit antiviral immune functions.

**Figure 4.**
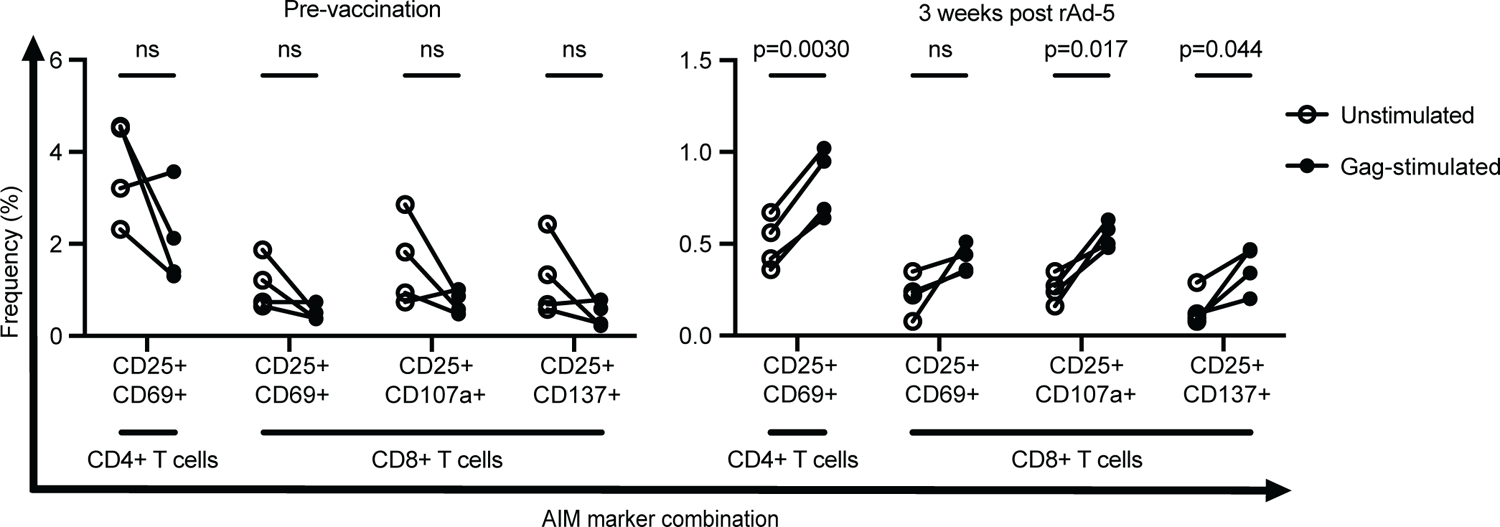
Vaccination enhances the frequency of CD4+ and CD8+ T cells in the PBMC that express activation-induced markers in response to Gag stimulation. The frequency of CD4+ T cells double-positive for CD25/CD69, and CD8+ T cells double-positive for CD25/CD69, CD25/ CD107a, and CD25/CD137 are plotted from time points before vaccination (left) and 3 weeks after the final vaccination (right). Frequencies are compared between unstimulated conditions (open circles) and stimulation with a Gag peptide pool (closed circles) in vitro. *P* values were calculated using paired T-tests.

### N-803 treatment is associated with increased Ki-67+ SIV-specific CD8+ T cells and continued T_EM_ bias of vaccine-stimulated CD8+ T cells

We wanted to determine whether N-803 could enhance SIV-specific vaccine-elicited T cells when compared to those naturally elicited by infection. ART was discontinued in all eight animals six weeks after the last vaccine or saline administration. The four vaccinated animals, but not the control animals, received three subcutaneous 0.1mg/kg doses of N-803, separated by two weeks each, beginning three days after ART cessation (Figure 1A). We used Gag_386-394_GW9 and Nef_103-111_RM9 tetramers to compare vaccine-elicited to naturally elicited CD8+ T cells, respectively. The Nef_103-111_RM9 tetramer+ CD8+ T cells (henceforth referred to as Nef RM9-specific CD8+ T cells) could only be elicited by SIV infection, as Nef was not included in the vaccine. One clear caveat is that some Gag GW9-specific CD8+ T cells were also elicited by SIV infection, in addition to the vaccine-induced Gag-responsive cells. Nonetheless, distinguishing between Gag GW9- and Nef RM9-specific CD8+ T cells represents one way to compare vaccine-elicited and naturally elicited CD8+ T cells within the same animals.

In the peripheral blood, the frequency of Gag GW9-specific CD8+ T cells was generally higher than Nef RM9-specific CD8+ T cells following vaccination and ART release, and throughout N-803 treatment (Figure 5A), as measured by the total area under the curve (AUC) for the two populations throughout N-803 treatment. The AUC for the Ki-67+ Gag GW9-specific CD8+ T cells was similar to the AUC for the Ki-67+ Nef RM9-specific CD8+ T cells (Figure 5B). This suggests that N-803 enhances the proliferation of antigen-specific CD8+ T cells indiscriminately whether those cells were elicited by vaccination or by natural infection. We then compared the total AUC of the frequencies of the Gag GW9- and Nef RM9-specific CD8+ T_CM_ and T_EM_ cells (gated on CD28+CD95+ or CD28-CD95+ cells within the tetramer+ parent population, respectively). Although some time points had to be excluded due to low numbers of cells, the AUC of Nef RM9-specific CD8+ T_CM_ cells was higher than Gag GW9-specific CD8+ T_CM_ cells (Figure 5C) and the AUC of Gag GW9-specific CD8+ T_EM_ cells was higher than the Nef RM9-specific CD8+ T_EM_ cells (Figure 5D). While in general the Gag-specific CD8+ T cells remained T_EM_ and the Nef-specific CD8+ T cells remained T_CM_, we detected no N-803-mediated changes in Gag- or Nef-specific CD8+ T cell frequencies or the distribution of T_CM_ or T_EM_ phenotypes. N-803 also transiently increased the number and frequency of bulk, Ki-67+, CD69+, and Ki-67+CD69+ CD4+ T cells, CD8+ T cells, and CD4+ and CD8+ T_CM_, T_TM_, and T_EM_ populations in the peripheral blood of the vaccinated animals (Supplemental Figures 3 and 4).

**Figure 5.**
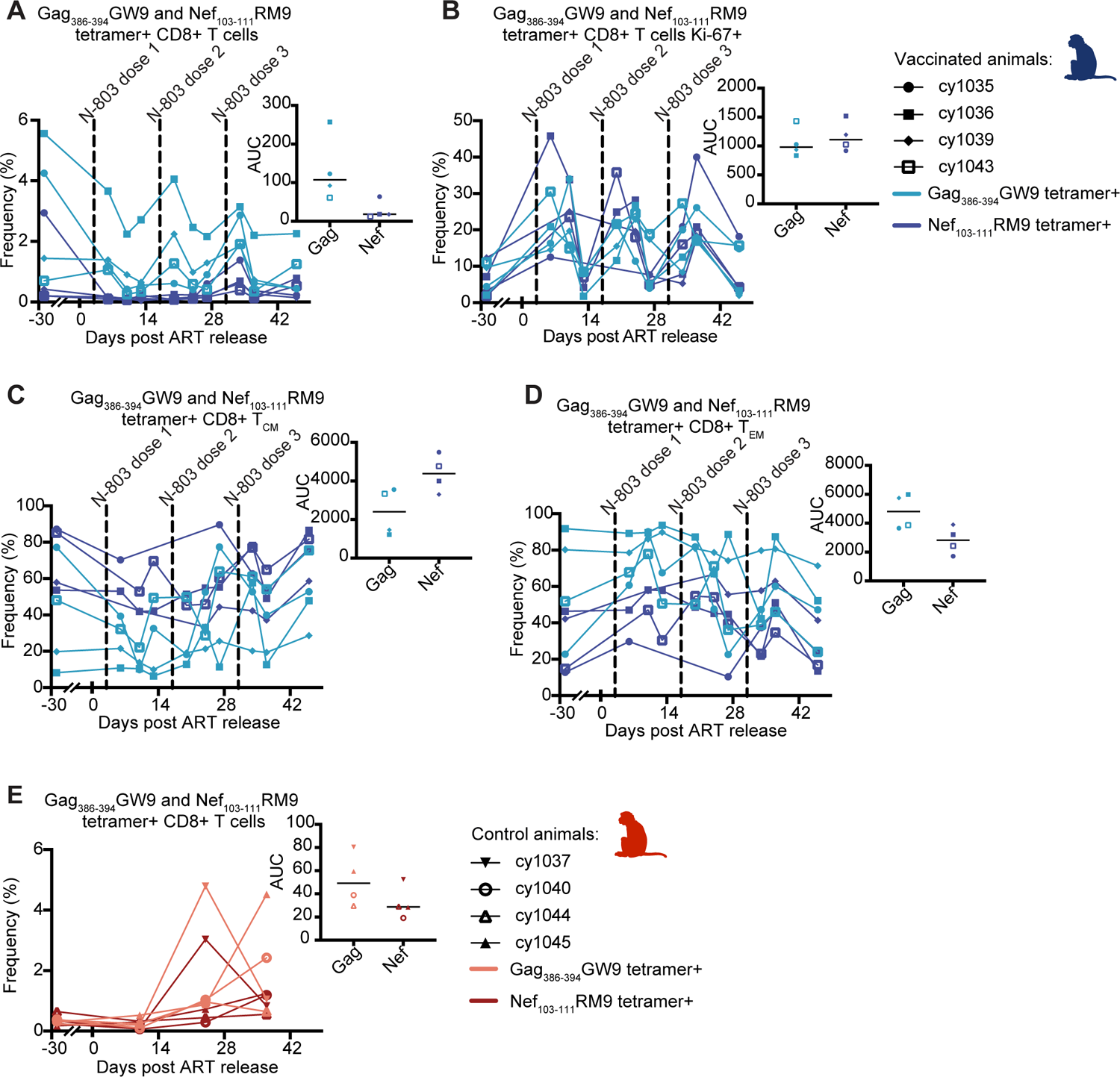
Changes in Gag GW9- and Nef RM9-specific CD8+ T cells in the PBMC of the vaccinated animals (blue tones) or control animals (red tones) following ART removal. **(A)** Frequency and AUC of Gag GW9-specific CD8+ T cells (light blue) compared to Nef RM9-specific CD8+ T cells (dark blue) (gated on tetramer+ cells within the parent CD8+ population) throughout the N-803 phase in the vaccinated animals. **(B)** Frequency and AUC of proliferating (Ki-67+) Gag GW9- or Nef RM9-specific CD8+ T cells throughout the N-803 phase in the vaccinated animals (gated on Ki-67+ cells within the tetramer+ parent population). **(C)** Frequency and AUC of Gag GW9-compared to Nef RM9-specific CD8+ T_CM_ (gated on CD28+CD95+ cells within the tetramer+ parent population) throughout the N-803 phase. **(D)** Frequency and AUC of Gag GW9-compared to Nef RM9-specific CD8+ T_EM_ (gated on CD28-CD95+ cells within the tetramer+ parent population) throughout the N-803 phase. **(E)** Frequency and AUC of Gag GW9-specific CD8+ T cells (light red) compared to Nef RM9-specific CD8+ T cells (dark red) (gated on tetramer+ cells within the parent CD8+ population) in the control animals following ART release. Results are displayed for each animal individually.

In the control animals, the Gag GW9- and Nef RM9-specific CD8+ T cells (Figure 5E) remained at similar frequencies prior to and following ART release and exhibited comparable AUCs. There was a slight increase in the frequency of these populations ∼three weeks after ART release. This was likely attributed to increased antigen exposure when ART was no longer present to suppress virus replication. Small sample sizes precluded statistical analyses of these paired data.

### Gag GW9- and Nef RM9-specific CD8+ T cell frequencies in the LN are similar during N-803

Because N-803 treatment can cause migration of CD8+ T cells to the LN (44, 45), we investigated whether vaccine-elicited T cells were found at higher frequencies than naturally-elicited T cells in the LN. We again used Gag_386-394_GW9 and Nef_103-111_RM9 tetramers to measure the phenotypes and frequencies of these vaccine- and naturally elicited populations of antigen-specific T cells, respectively.

In the LNs of the treatment animals, the frequency of Gag GW9-specific CD8+ cells (range of 0.11-5.13%) was up to 10-fold higher than the frequency of Nef RM9-specific CD8+ T cells (range of 0.11-0.54%), and the frequencies of bulk Gag- and Nef-specific CD8+ T cells in the LNs were unaffected by N-803 treatment (Figure 6A). The Gag GW9- and Nef RM9-specific T_CM_ and T_EM_ in the LN of the treatment animals were also unchanged throughout the N-803 phase. Like in the peripheral blood, the frequency of Gag GW9-specific T_CM_ was lower than Nef RM9-specific T_CM_, and the frequency of Gag GW9-specific T_EM_ was higher than Nef RM9-specific T_EM_ cells in the LN (Figures 6B and C).

**Figure 6.**
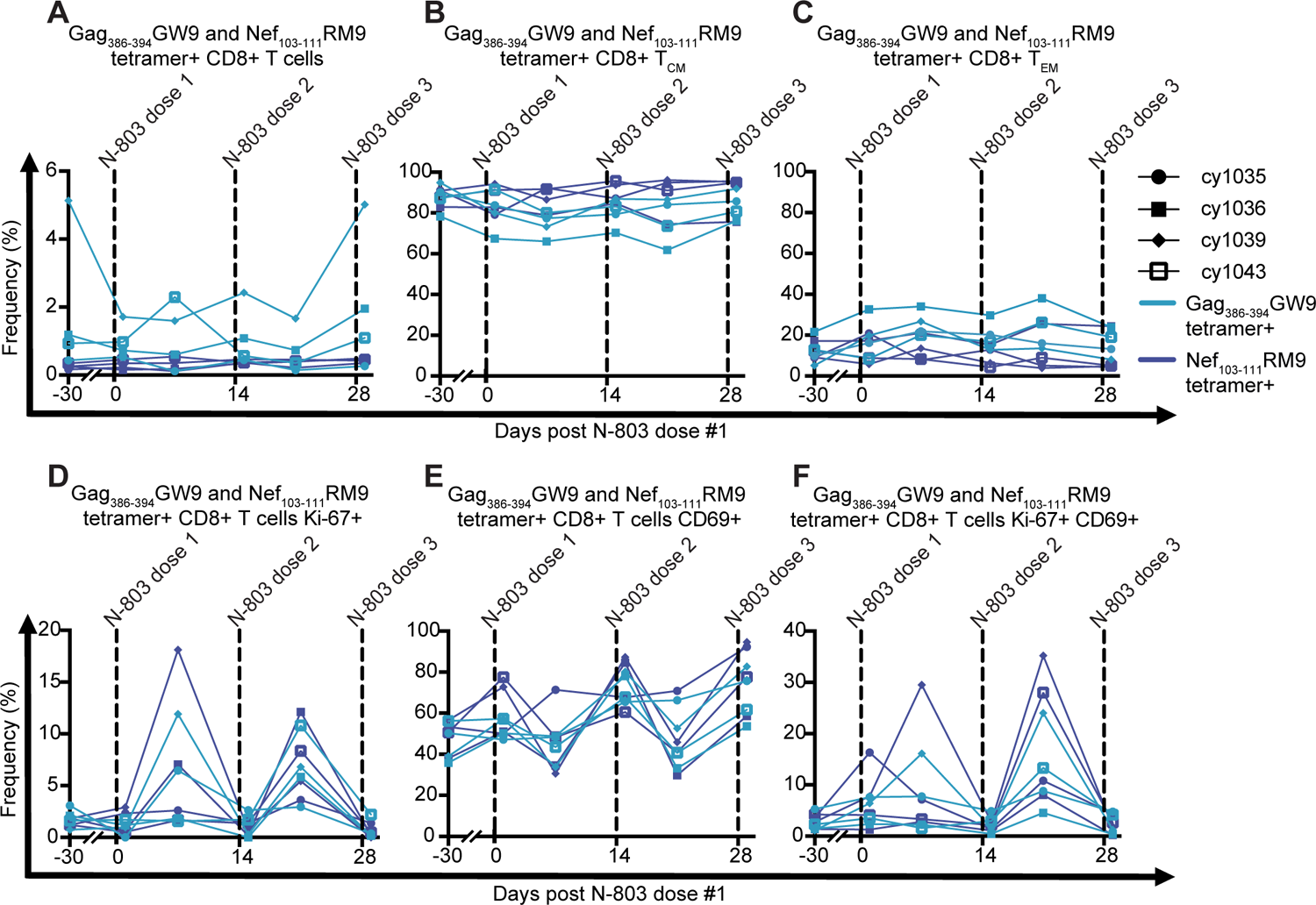
N-803-mediated changes in Gag GW9- and Nef RM9-specific CD8+ T cells in the LN of the vaccinated animals. **(A)** Frequency of Gag GW9-specific CD8+ T cells (light blue) compared to Nef RM9-specific CD8+ T cells (dark blue) (gated on tetramer+ cells within the parent CD8+ population) throughout the N-803 phase. **(B)** Frequency of Gag GW9-compared to Nef RM9-specific CD8+ T_CM_ (gated on CD28+CD95+ cells within the tetramer+ parent population) throughout the N-803 phase. **(C)** Frequency of Gag GW9-compared to Nef RM9-specific CD8+ T_EM_ (gated on CD28-CD95+ cells within the tetramer+ parent population) throughout the N-803 phase. **(D-F)** Frequency of Gag GW9- or Nef RM9-specific CD8+ T cells expressing the proliferation marker Ki-67 alone **(D)**, the activation marker CD69 alone **(E)**, or Ki-67 and CD69 together **(F)** throughout the N-803 phase, each gated within the parent tetramer+ population. Results are displayed for each animal individually.

We then examined the frequency of Gag GW9- and Nef RM9-specific CD8+ T cells expressing either Ki-67, CD69, or both Ki-67 and CD69 in the LN. The effect of N-803 on the subpopulations of each of these tetramer+ parent populations was similar (Figure 6D-F). Thus, in the LN, vaccine- and naturally elicited antigen-specific CD8+ T cells responded similarly to N-803 treatment. Unfortunately, LNs could not be collected 7 days after the third dose, so we were unable to evaluate whether the proliferative responses were enhanced to a higher degree after the third N-803 dose. The fold change of bulk CD4+ and CD8+ T cells, as well as the frequency of CD4+ and CD8+ T_CM_, T_TM_, and T_EM_ expressing Ki-67 and/or CD69 in the LN typically increased in the vaccinated animals after each of the first two doses of N-803, mirroring the results of the Gag GW9- and Nef RM9-specific populations in the LN (Supplemental Figure 5). As expected, the bulk populations of T cells were unaffected in the control animals over time.

### SIV was not consistently detected in the plasma during N-803 treatment

While N-803 treatment in vivo increased the number, frequency, activation, and proliferation of CD4+ and CD8+ T cell subsets (Supplemental Figures 2-4), these cellular changes had no apparent effect on SIV plasma viremia, which remained below 10^4^ copies/mL following ART discontinuation. Up to 60 days after suspending ART, one vaccinated animal (cy1035) and two control animals (cy1037 and cy1040) had transient, detectable low-level viremia (between 1.2×10^2^ and 6×10^3^ceq/mL) (Figure 7A). Viremia was undetectable in all animals 60 days after ART release. Further studies to understand the lack of rebound in both groups of animals following ART release are underway (manuscript in preparation). There was no significant difference in the AUCs of the viral loads between the vaccinated and control animals during the 60-day period following ART release evaluated in this study (Figure 7B).

**Figure 7.**
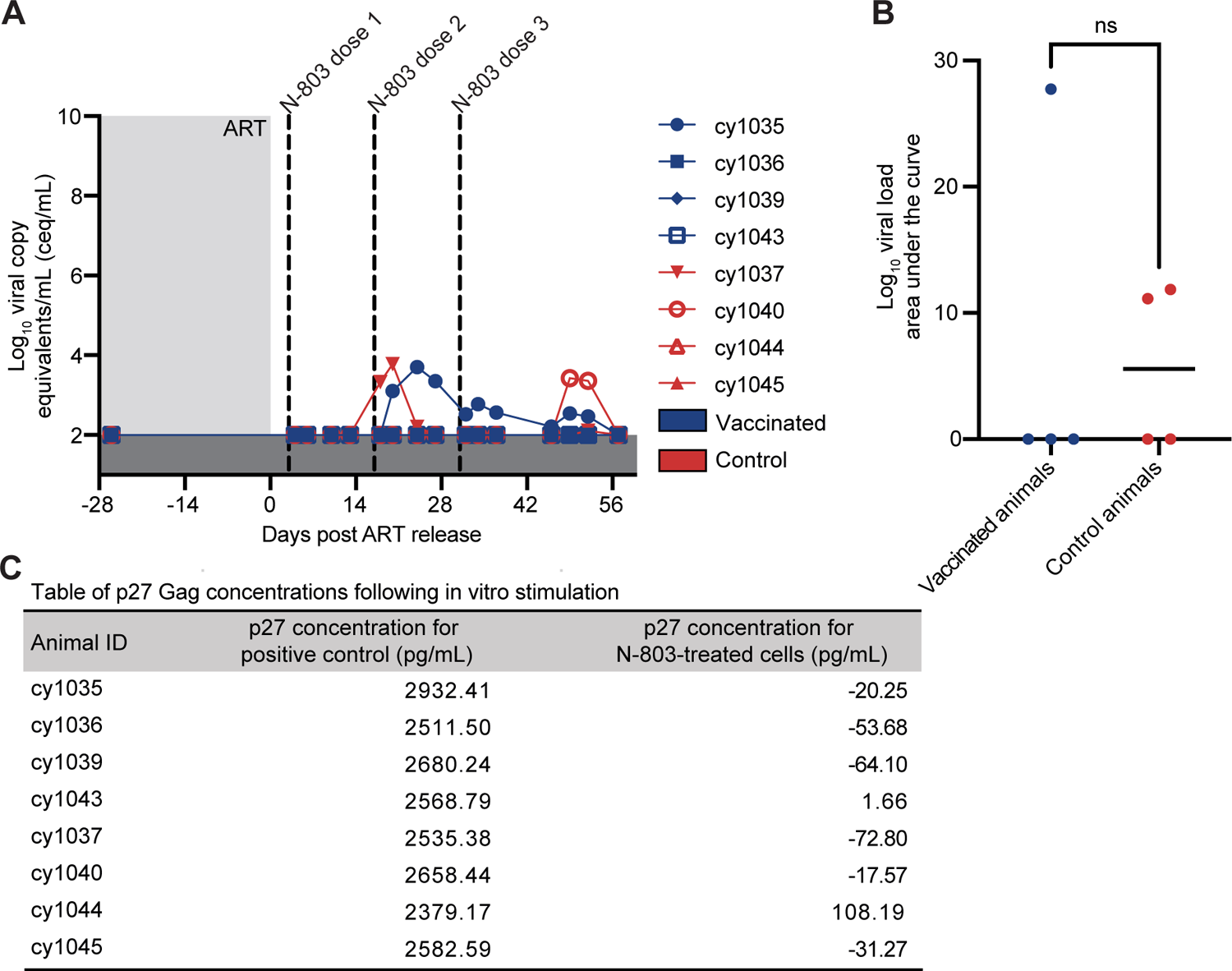
Detection of SIV in vivo and in vitro following N-803 treatment. **(A)** Plasma viral loads following ART release: viral loads are displayed as log10ceq/mL. **(B)** Log_10_viral load AUC analysis from day 0-60 post ART release. Results are displayed as median and individual values. *P* value was calculated using a Mann-Whitney U test. **(C)** Table of supernatant p27 concentrations from CD4+ T cells treated with anti-CD3/CD28 beads (center column) or N-803 (right column).

We also treated isolated CD4+ T cells from both groups of animals with N-803 in vitro. Consistent with our in vivo observations, N-803 did not induce the production of p27 Gag (Figure 7C). As a positive control, treatment of cells with anti-CD3/anti-CD28 beads readily induced virus production, leading to p27 Gag concentrations ranging from 2379.17 to 2932.41pg/mL in the supernatants (Figure 7C). We thus observed no direct increase in SIV replication due to N-803 in vivo or in vitro, despite profound N-803-mediated changes in CD4+ and CD8+ lymphocyte populations and an absence of ART.

## Discussion

Here, we evaluated an SIV therapeutic vaccine strategy using heterologous viral vectors delivering Gag immunogens to SIV+ MCMs receiving ART, followed by immunotherapy with N-803 after ART discontinuation, as a novel therapeutic regimen to induce anti-SIV cellular immunity. We hypothesized that this regimen would generate and recall Gag-specific CD8+ T cells. To our knowledge, this is the first study to evaluate the combination of therapeutic i.v. HPBB vaccine administration followed by N-803 treatment after suspending ART. We found that this vaccine regimen elicited a high magnitude of Gag-specific CD8+ T cells that contracted following each vaccine dose. These vaccine-elicited Gag-specific cells were boosted by N-803 to a similar degree as naturally elicited Nef-specific CD8+ T cells (Figures 5 and 6). Because durable rebound was not detected in any animal up to 60 days post-ART cessation, it remains unclear whether therapeutically elicited immune responses would have been sufficient to suppress SIV replication. This lack of rebound will be investigated in further studies.

One goal of the vaccine regimen presented in this study was to elicit broad and abundant Gag-specific CD8+ T cells. Broader responses to Gag have been associated with strong antiviral activity and decreased viral loads (8, 31, 32). High magnitude Gag-specific CD8+ T cells have been associated with CD4+ T cell preservation and lower HIV viral loads (31). Our vaccine regimen modestly increased the breadth of Gag-specific T cells targeting Gag_386-394_GW9 and four additional Gag epitopes (Gag_459-467_TV9, Gag_146-154_HL9, Gag_28-37_KA10, and Gag_221-229_PR9) when compared to the unvaccinated animals. After vaccination, there was also an increase in the frequency of Gag-specific CD4+ and CD8+ T cells expressing AIM markers, suggesting that vaccination enhanced the frequency of cells able to recognize and respond to Gag by way of antigen recognition, activation, and degranulation.

Our vaccine regimen transiently increased the frequency of Gag GW9-specific CD8+ T cells in the PBMC. These cells had a T_EM_ phenotype and expressed the proliferation marker Ki-67. Whereas both longer and shorter intervals (2 weeks to multiple months between doses) can elicit high frequencies of CD8+ T cells with a similar potential to proliferate and respond to antigen re-exposure, very short intervals (2 weeks) elicit CD8+ T cell populations that are more susceptible to contraction in mice (20). Petitdemange et al. found that proliferating Gag-specific CD8+ T cells elicited by HPBB vaccination delivered to macaques in longer intervals (weeks 1, 9, and 37) expanded to a higher, more sustained, frequency after the final boost compared to our HPBB vaccine regimen, which was given in four-week intervals (15). While Gag GW9-specific CD8+ T cells did expand in our study, they subsequently contracted to nearly pre-vaccine levels after each dose (Figure 2). The differences in magnitude and persistence of Gag-specific CD8+ T cells that we observed and those detected in the Petitdemange study could stem from a variety of sources. First, an HPBB vaccine regimen may require longer intervals than we used to elicit longevous cell populations. Second, we used a non-replicating MVA vector for the second immunization and Petitdemange et al. used a replicating vaccinia virus (VV) vector. The inclusion of a more pathogenic VV vector rather than the attenuated MVA vector may have improved the efficacy of this boost. Finally, although our animals were receiving ART, the initial insult of SIV infection and underlying impact on the host cellular immunity could weaken the responsiveness to vaccination. It would be interesting to evaluate this HPBB vaccine regimen in SIV-naïve MCMs to determine if it elicits a persistently higher frequency of GW9-specific CD8+ T cells.

Because all the vaccines delivered SIVmac239 Gag in this study, we were able to use Gag_386-394_GW9 and Nef_103-111_RM9 tetramers to compare vaccine-elicited (Gag GW9-specific) to naturally elicited (Nef RM9-specific) CD8+ T cells within each animal. Many therapeutic vaccine regimens deliver multiple SIV immunogens, such as Gag, Pol, and Env (23, 29, 30), making any distinction between vaccine-elicited cells and cells naturally elicited by HIV/SIV infection difficult. Prophylactic vaccine studies delivering multiple immunogens similarly lose the ability to distinguish between antigen-specific CD8+ T cells elicited by vaccination or SIV infection (57, 58). Further, there is limited availability of tetrameric reagents to track defined antigen-specific CD8+ T cells in individuals even when their MHC genetics are known (17, 45, 59). These peptide-loaded MHC class I tetramers are required to distinguish between CD8+ T cells targeting immunogens present in the vaccine or challenge virus. By using MCMs expressing the *Mafa-A1*063* MHC class I molecule that restricts the Gag_386-394_GW9 epitope in the vaccine immunogen and the Nef_103-111_RM9 epitope in the SIV challenge virus, we could distinguish between cells elicited by vaccination versus SIV infection within the same animals.

As N-803 has been shown to enhance the frequency and LN trafficking of SIV-specific CD8+ T cells elicited by SIV infection (44, 45), we tested the hypothesis that N-803 would boost the magnitude of vaccine-elicited CD8+ T cells to a higher degree than those CD8+ T cells elicited naturally by SIV infection. However, we actually found that N-803 treatment similarly increased the frequency of Ki-67+ Gag GW9- and Nef RM9-specific CD8+ T cells in the PBMC, suggesting a role for N-803 in the proliferation of all antigen-specific CD8+ T cells. This is consistent with defined contributions of IL-15 in CD4+ and CD8+ T cell activation, proliferation, and maintenance (60, 61). Notably, since the vaccine regimen elicited a high frequency of Gag GW9-specific CD8+ T cells, N-803 treatment further magnified the frequency of CD8+ T cells elicited by the vaccine compared to those elicited by natural infection. In the context of antiviral immunity, the potential of N-803 to drive activation and proliferation of vaccine-elicited CD8+ T cells may support the use of N-803 as an adjuvant in HIV/SIV therapeutic strategies.

We were surprised that N-803 treatment did not dramatically increase the frequency of Gag GW9 specific CD8+ T cells in the LNs. It is possible that N-803 directed these antigen-specific CD8+ T cells to different tissue sites like the gut, which we did not sample. Alternatively, the presence of antigen may be required for N-803-mediated LN localization of antigen-specific T cells. As only one vaccinated animal exhibited low-level plasma viremia and the other three vaccinated animals were aviremic at the time of N-803 administration, it is unclear whether higher viremia and therefore higher levels of antigen-stimulation would have enabled N-803 to direct more antigen-specific T cells to the LNs. It would be interesting to deliver N-803 and vaccines concomitantly to determine if combining N-803 with antigen stimulation would enhance the magnitude of SIV-specific CD8+ T cells directed to the LNs.

Antigen-specific CD8+ T_EM_ are critical for HIV/SIV therapeutic strategies because lymphocytes differentiated into T_EM_ phenotypes have been associated with enhanced effector function, progressive clearance of SIV reservoirs, and detection of replication-competent virus in resting CD4+ T cells (62–64). The vaccinated animals here developed a higher frequency of Gag GW9-specific T_EM_ than the control animals. The vaccinated animals also developed a higher frequency of Gag GW9-specific CD8+ T_EM_ than Gag GW9-specific T_CM_, consistent with reports that replicating vectors like VSV provide recurrent antigen exposure that favor the induction of protective T_EM_ responses (27). This represents one potentially advantageous feature of this vaccine regimen. Interestingly, Hansen and colleagues found that protective SIV-specific T_EM_ populations elicited by persistent rhesus cytomegalovirus (RhCMV) vectors prior to SIV infection may have played a role in preventing SIV infection (62). In our study, the T_EM_ bias of vaccine-elicited Gag-specific CD8+ T cells and T_CM_ bias of Nef-specific CD8+ T cells elicited by SIV infection were maintained and generally unaltered throughout N-803 treatment. Whether N-803 would boost T_EM_ cells in the case of rhCMV vaccination strategy, or whether the HPBB vaccine strategy described here combined with N-803 treatment would be protective if delivered prophylactically remains unclear. However, we propose that alternative ways exist to elicit and expand antigen-specific CD8+ T_EM_ cells that do not require vaccination with a variant strain of CMV.

One drawback of this study is the lack of additional single-intervention animal cohorts. Ideally, the inclusion of one cohort treated with the vaccine regimen but not N-803 and another cohort treated with N-803 and no vaccinations would have enabled a more thorough evaluation of the immune responses of each intervention individually. These single intervention cohorts would allow us to define how vaccine-elicited CD8+ T cells respond to ART interruption in the absence of additional cytokine modulation, and how N-803 boosts lymphocyte populations untouched by a therapeutic vaccine intervention when delivered after ART release. However, macaques are a valuable and costly resource. Thus, rather than add two additional cohorts of animals, we evaluated the impact of N-803 on vaccine-elicited CD8+ T cells versus CD8+ T cells elicited only by SIV infection within the vaccinated group.

It was unexpected that N-803 treatment, alone, did not induce SIV replication in vivo or in vitro. The role of N-803 as a latency-reversing agent (LRA) remains unclear, despite in vitro and in vivo evaluation (45, 65–67). Jones et al. described N-803 treatment of cell cultures in vitro induced antigen presentation and CD8+ T cell recognition in primary cell cultures and ex vivo cultures from HIV+ individuals (65). McBrien and colleagues found that N-803 reactivated HIV expression from CD4+ T cells infected in vitro, but that reactivation could be inhibited by the presence of activated CD8+ T cells (67). These results stand in contrast to our observations that in vitro N-803 treatment of CD4+ T cells isolated from the PBMC of SIV+ MCM did not induce p27 Gag expression. It is possible that our p27 ELISA assay was not sensitive enough to detect very low levels of N-803-mediated latency reversal in vitro. Webb et al. also show that in vivo N-803 administration was insufficient to perturb the viral reservoir (45). Thus, it was not surprising that N-803 did not induce viremia in vivo after ART interruption of MCM with an intact CD8 compartment, but the failure of N-803 to induce p27 expression in vitro suggests that this agent did not perturb the reservoir. It may therefore be necessary to combine N-803 with a different LRA for a “shock and kill” strategy to effectively reduce the size of the viral reservoir.

In conclusion, we describe the immunogenicity and immunomodulation of a novel therapeutic regimen combining HPBB vaccination delivering Gag immunogens during ART combined with N-803 after suspending ART. The vaccine regimen elicited a high magnitude of proliferative and activated CD8+ T cells driven toward a T_EM_ phenotype. N-803 enhanced both the vaccine-elicited and naturally elicited SIV-specific CD8+ cells. Thus, N-803 remains an attractive immunomodulatory agent for HIV/SIV to enhance cellular immune responses elicited both by infection and by other therapeutic interventions. Any functional cure for HIV will surely involve a combination of multiple therapeutic interventions, and we show here that N-803 combined with vaccination generates and recalls cellular immune responses against SIV.

## Materials and Methods

### Animal care and use

Eight male Mauritian cynomolgus macaques (MCMs) were purchased from Bioculture Ltd. and were housed and cared for by the Wisconsin National Primate Research Center (WNPRC) according to protocols approved by the University of Wisconsin Graduate School Animal Care and Use Committee (IACUC; protocol number G005507). The animals were chosen based on the presence of at least one copy of the M3 MHC haplotype and the absence of the M1 haplotype associated with viral control (Table 1) (46, 47, 68). All eight MCMs were infected intravenously (i.v.) with 10,000 IUs of SIVmac239M (69). An antiretroviral therapy (ART) regimen consisting of 2.5mg/kg dolutegravir (DTG, from ViiV Healthcare, Research Triangle, NC), 5.1mg/kg tenofovir disoproxil fumarate (TDF, from Gilead, Foster City, CA), and 40mg/kg emtricitabine (FTC, from Gilead) in 15% Kleptose (Roquette America) in water was delivered subcutaneously daily beginning on day 14 post-infection. Four animals (the vaccinated group) received Gag or Gag-Pol proteins encoded by three heterologous viral vectors. Recombinant vesicular stomatitis virus New Jersey strain (VSV) expressed full-length SIVmac239 Gag (58, 70). The VSV-Gag vector was given i.v. at a dose of 5×10^7^ PFU per animal at week 23 post-SIV. Recombinant modified vaccinia virus Ankara strain (MVA) expressed SIVmac239 Gag-Pol (71). The MVA-Gag-Pol vector was given i.v. at a dose of 1×10^8^ PFU per animal at week 27 post-SIV. Recombinant adenovirus serotype 5 (rAd-5) expressed full-length SIVmac239 Gag (ViraQuest Inc.). The rAd-5-Gag was given i.v. at a dose of 7×10^10^ particles per animal at week 31 post-SIV. All immunogens were injected i.v. in 1mL total volume, diluted in sterile 1xPBS, where necessary. Four animals (the control group) received 1mL sterile 1xPBS injections only. Three doses of N-803, separated by 14 days each, were delivered subcutaneously to the four treatment animals at a dose of 0.1mg/kg beginning three days after ART interruption.

### Plasma viral load analysis

Plasma was isolated from undiluted whole blood by Ficoll-based density centrifugation and cryopreserved at −80°C. Plasma viral loads were quantified as previously described (72). Briefly, the Maxwell Viral Total Nucleic Acid Purification kit (Promega, Madison, WI) was used to isolate viral RNA (vRNA) from plasma samples. vRNA was then reverse transcribed using the TaqMan Fast Virus 1-Step qRT-PCR kit (Invitrogen) and quantified on a LightCycler 480 (Roche, Indianapolis, IN).

### IFN-γ ELISPOT assays

IFN-γ ELISPOT assays were performed using fresh and cryopreserved PBMC, as previously described (73). Peptides (Gag_386-394_GW9, Nef_103-111_RM9, Gag_459-467_TV9, Gag_146-154_HL9, Gag_28-37_KA10, and Gag_221-229_PR9, and a Gag peptide pool containing 15-mer peptides spanning the full SIVmac239 Gag proteome, each overlapping by 11 amino acids [NIH HIV Reagent Program, managed by ATCC]) were selected from epitopes known to be restricted by the *Mafa-A1*063* MHC class I allele expressed on the M3 MHC haplotype (50). PBMC were isolated from EDTA-anticoagulated blood by Ficoll-based density centrifugation. Precoated monkey IFN-γ ELISPOTplus plates (Mabtech, Cincinnati, OH) were blocked with R10 (RPMI 1640 supplemented with 10% FBS, 1% antibiotic-antimycotic [Thermo Fisher Scientific, Waltham, MA], and 1% L-glutamine [Thermo Fisher Scientific]), and individual peptides were added to each well at a final concentration of 10μM. The Gag peptide pool was added to cells at a final concentration of 625μg/mL (5μg/mL of each peptide). Each peptide or peptide pool was tested in duplicate. Concanavalin A (10μM) was used as a positive control and was tested in duplicate as well. Four wells per animal received no peptides as a negative control to calculate background reactivity. Plates were incubated overnight at 37°C in 5% CO_2_. Assays were performed according to the manufacturer’s protocol, and wells were imaged with an ELISPOT plate reader (AID Autoimmun Diagnostika GmbH). Positive responses were determined using a one-tailed t-test at an α level of 0.05, where the null hypothesis was that the background level would be greater than or equal to the treatment level (57, 73). Statistically positive responses were considered valid only if both duplicate wells contained 50 or more spot-forming cells (SFCs) per 10^6^ PBMC. If statistically positive and ≥50 SFCs per 10^6^ PBMC, the reported values are the average of the two test wells minus the average of all four negative control wells.

### Tetramerization of Gag_386-394_GW9 and Nef_103-111_RM9

The Gag_386-394_GW9 and Nef_103-111_RM9 peptides were purchased from Genscript (Piscataway, NJ). The NIH Tetramer Core Facility at Emory University (Atlanta, GA) produced biotinylated Mafa-A1*063 MHC class I monomers loaded with these peptides. The Mafa-A1*063 Gag_386-394_GW9 and Mafa-A1*063 Nef_103-111_RM9 monomers were tetramerized with streptavidin-PE (0.5mg/mL, BD biosciences) and streptavidin-BV421 (0.1mg/mL, BD biosciences), respectively, at a 4:1 molar ratio of monomer:streptavidin in the presence of a 1x protease inhibitor cocktail solution (Calbiochem, Millipore Sigma). 1/5^th^ volumes of streptavidin-PE or streptavidin-BV421 were added to each monomer every 20 minutes and incubated, rotating in the dark at 4°C until the full streptavidin volume was added.

### Phenotype staining of T cells by flow cytometry

Previously frozen PBMC isolated from whole blood and previously frozen lymph node (LN) mononuclear cells (LNMC) isolated from LN biopsies were used to assess the quantity and phenotype of T cell populations longitudinally. Briefly, cells were thawed, washed once with R10, and rested for 30 minutes at room temperature in a buffer consisting of 2% FBS in 1X PBS (2% FACS buffer) with 50nM dasatinib (Thermo Fisher Scientific). Cells were washed once with 2% FACS buffer with 50nM dasatinib and incubated with the Gag_386-394_GW9 and Nef_103-111_RM9 tetramers for 45 minutes at room temperature. Cells were then washed once with 2% FACS buffer with 50nM dasatinib and incubated with the remaining surface markers (Table 2) for 20 minutes at room temperature. Cells were washed twice with 2% FACS buffer with 50nM dasatinib and fixed using fixation/permeabilization solution (Cytofix/Cytoperm™ fixation and permeabilization kit, BD Biosciences) for 20 minutes at 4°C. Cells were next washed twice with cold 1x Perm/Wash^TM^ buffer (Cytofix/Cytoperm™ fixation and permeabilization kit, BD Biosciences) and incubated with a master mix containing 95μL of 1x Perm/Wash^TM^ buffer and 5μL of the intracellular marker Ki-67 (Table 2) for 20 minutes at 4°C. Cells were then washed twice with 1x Perm/Wash^TM^ buffer and acquired immediately using a FACS Symphony A3 (BD Biosciences). The data were analyzed using FlowJo software for Macintosh (BD Biosciences, version 10.8.0). Subpopulations of cells were excluded from analysis when the parent population contained <50 events.

**TABLE 2.** Antibodies used for T cell phenotypinig

| **Antibody** | **Clone** | **Fluorochrome** | **Surface or intracellular** |
| --- | --- | --- | --- |
| Live/Dead |  | Near-infrared | Surface |
| CD3 | SP34-2 | BUV563 | Surface |
| CD4 | SK3 | BUV737 | Surface |
| CD8 | RPA-T8 | BUV395 | Surface |
| CD28 | CD28.2 | BUV661 | Surface |
| CD69 | TP1.55.3 | PE-Texas Red (ECD) | Surface |
| CD95 | DX2 | BV711 | Surface |
| CCR7 | 150503 | FITC | Surface |
| CXCR5 | MU5UBEE | PE-Cy7 | Surface |
| Ki-67 | B56 | AF-647 | Intracellular |
| Gag GW9 tetramer | --- | PE | Surface |
| Nef RM9 tetramer | --- | BV421 | Surface |

### Activation-induced marker (AIM) assays

AIM assays were performed to characterize the antigen-specific markers of activation similarly to previously published work (51, 54, 74). Previously frozen PBMC isolated from whole blood and previously frozen LNMC isolated from LN biopsies were thawed, washed twice with R10, and incubated for ∼20 hours in Gibco AIM V^TM^ serum-free medium (Thermo Fisher Scientific) at 37°C in 5% CO_2_ with either AIM V^TM^ medium alone (unstimulated) or with a Gag peptide pool containing 15-mer peptides spanning the full SIVmac239 Gag proteome, each overlapping by 11 amino acids (provided by the HIV Reagent Program), at a final concentration of 62.5μg/mL (0.5μg/mL of each peptide) (Gag-stimulated). Two wells stimulated with 5μg/mL Concanavalin A (ConA) were included in each batch of staining as a positive control. Anti-CD107a and anti-CD154 antibodies (Table 3) were added to all cells during the stimulation. Following the stimulation, cells were washed twice with 2% FACS buffer and stained with antibodies to the indicated surface markers (Table 3) for 20 minutes at room temperature. Cells then were washed twice with 2% FACS buffer, fixed with 2% paraformaldehyde for 20 minutes at room temperature, washed twice more with 2% FACS buffer, and resuspended in 2% FACS buffer. Flow cytometry was performed as described above.

**TABLE 3.**
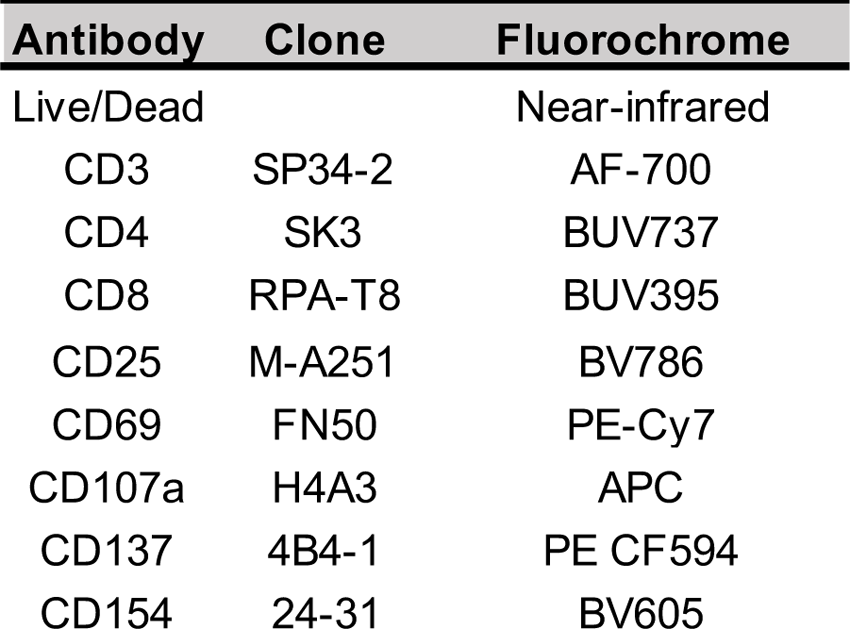
Antibodies used for AIM assay

### In vitro latency reactivation and p27 ELISA

Previously cryopreserved PBMC isolated from whole blood was thawed. After thawing, CD4+ T cells were isolated from the PBMC by negative selection using nonhuman primate CD4 MicroBeads according to the manufacturer’s protocol (Miltenyi Biotec). CD4+ T cells were incubated for 72 hours at 37°C with either anti-CD3/anti-CD28 beads (1:2 bead-to-cell ratio, Miltenyi Biotec) or 200ng/mL N-803 in the presence of 50µM raltegravir and 5µM saquinavir. After 72 hours, the supernatant was then collected, frozen, and subsequently subjected to SIV p27 ELISA per the manufacturer’s protocol (ZeptoMetrix). ELISA plates were immediately read using a GloMax®-Multi Detection System microplate reader (Promega) at 450nm absorbance.

### Statistical analysis

Area under the curve analyses were performed using GraphPad Prism. For statistical analyses in which animal groups were being compared to each other at the same time point, Mann-Whitney U tests were performed. Comparisons between unstimulated conditions and stimulated conditions within the same animal were tested using paired T-tests.

### Study Approval

This study was approved by the University of Wisconsin Graduate School Animal Care and Use Committee (IACUC; protocol number G005507).

## Author contributions

OEH, ALE, VV, PJS, and SLO contributed to the conception and design of the experiments. MRR, TCF, and SLO provided supervision and reviewed data. OEH, AJB, AJW, AMW, KNE, LMM, and AEG conducted experiments. OEH, ALE, LMM, and PTE analyzed the data. VV and JTS provided key reagents. OEH and SLO wrote the manuscript.

## Acknowledgments

The SIVmac239M was generously provided by Dr. Brandon Keele (Frederick National Laboratory for Cancer Research, Frederick, MD).

1. The DTG was graciously provided by ViiV Healthcare (Research Triangle, NC).
2. The TDF and FTC were graciously provided by Gilead (Foster City, CA).
3. The MVA was generously provided by Dr. Bernard Moss (NIH/NIAID).
4. The VSV was generously provided by Dr. Vaiva Vezys (University of Minnesota, Minneapolis, MN).
5. The N-803 was generously provided by ImmunityBio (Culver City, CA).
6. The following reagent was obtained through the NIH HIV Reagent Program, Division of AIDS, NIAID, NIH: Peptide Pool, Simian Immunodeficiency Virus (SIV)mac239 Gag Protein, ARP-12364, contributed by DAIDS/NIAID.
7. We thank the NIH Tetramer Core Facility (contract number 75N93020D00005) for generating the Mafa-A1*063 Gag_386-394_GW9 and Mafa-A1*063 Nef_103-111_RM9 biotinylated monomers.
8. We are grateful to the WNPRC staff for the exceptional veterinary care provided to the animals throughout this study.
9. The Wisconsin National Primate Research Center is supported by grants P51RR000167 and P51OD011106.
10. This study was funded through the National Institute of Health (NIH R01 AI108415).

---

**Supplemental Figure 1.**
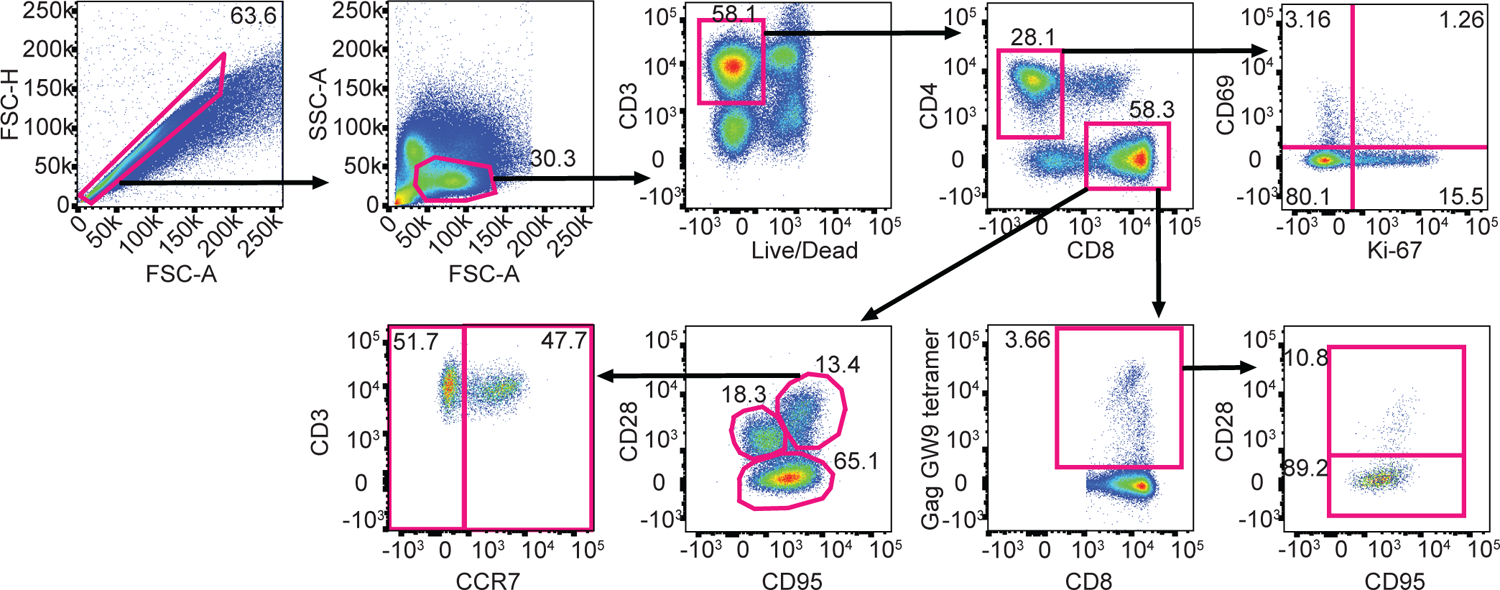
T cell gating schematic. Representative gating strategy used to evaluate memory phenotype, activation, and proliferation of bulk CD4+ and CD8+ T cells, and antigen-specific CD8+ T cells.

**Supplemental Figure 2.**
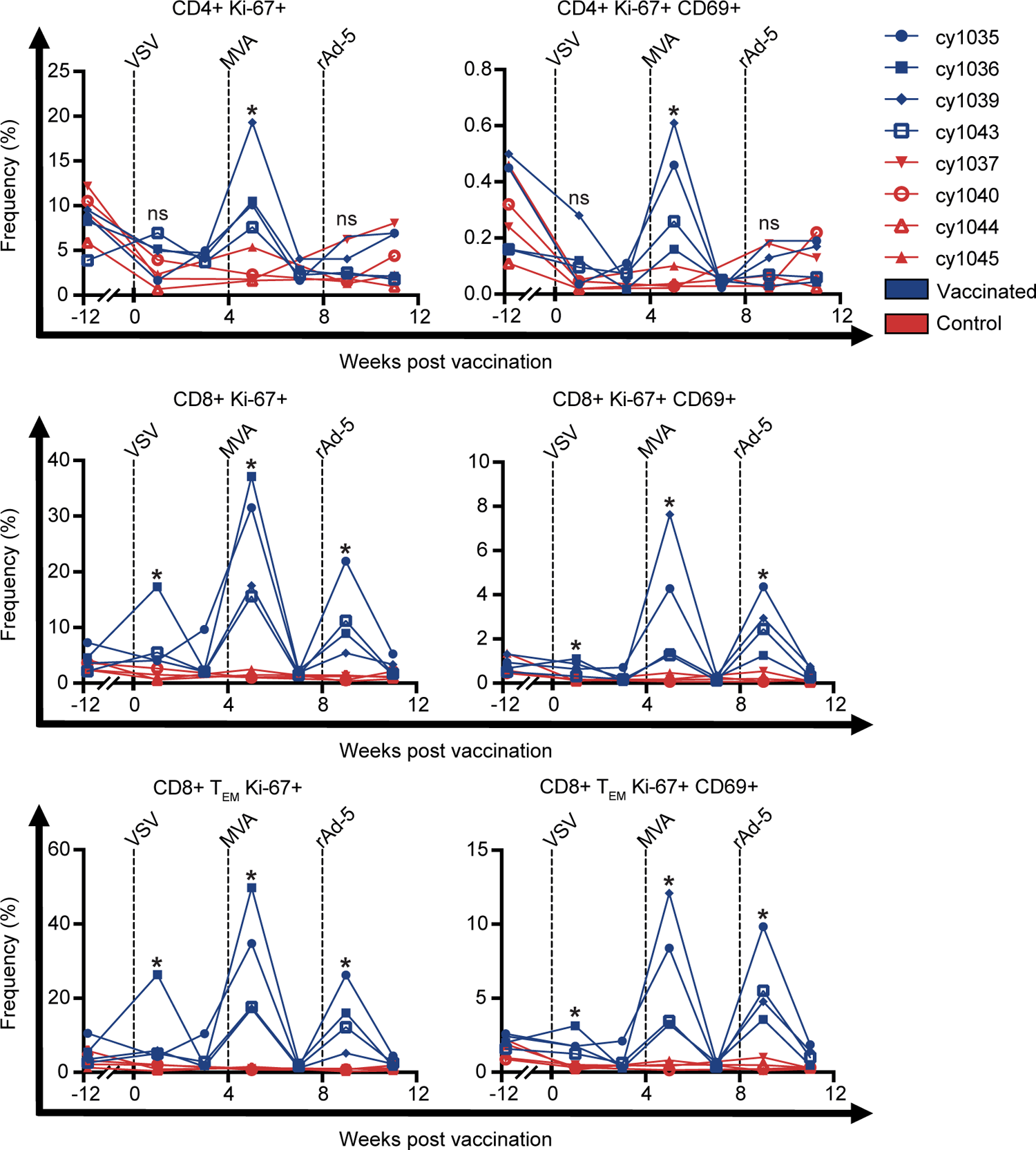
Proliferation and activation of CD4+ and CD8+ T cells in the PBMC throughout vaccination. Frequency of bulk CD4+ T cells (top row), bulk CD8+ T cells (middle row), and CD8+ T_EM_ (bottom row) in the PBMC expressing the proliferation marker Ki-67 alone (left) or in combination with the activation marker CD69 (right) throughout the vaccination phase. Results are displayed for each vaccinated (blue) and control (red) animal individually. * *P*=0.0286. *P* values were calculated using Mann-Whitney U tests comparing the vaccinated and control groups at each indicated point.

**Supplemental Figure 3.**
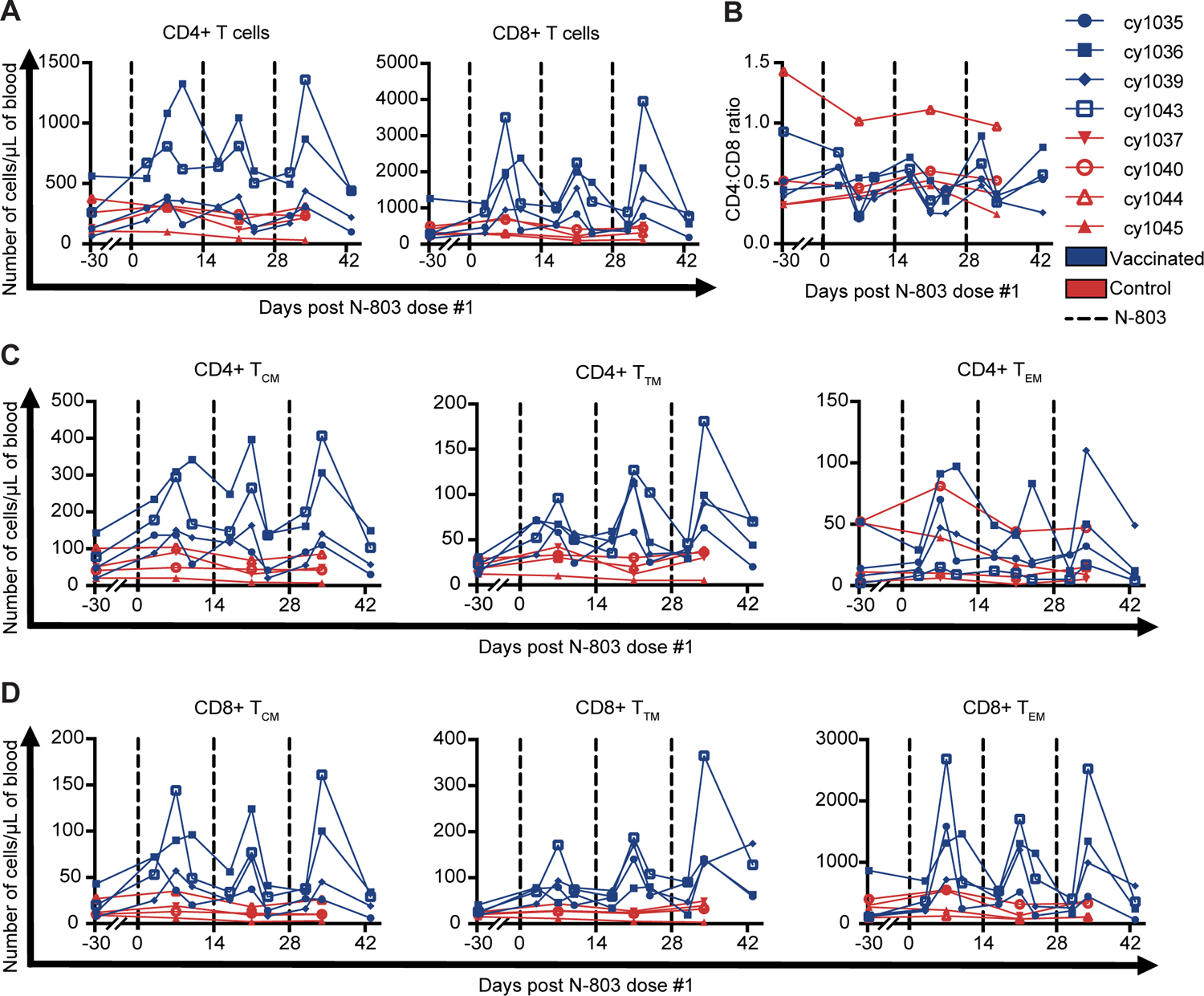
N-803-mediated changes in the absolute number of CD4+ T cell subsets, CD8+ T cell subsets, and CD4:CD8 ratio in the PBMC. (A) Number of CD4+ T cells (left) and CD8+ T cells (right) per μL of blood. **(B)** CD4:CD8 ratio. Number of CD4+ **(C)** and CD8+ **(D)** T_CM_ (left), T_TM_ (center), and T_EM_ (right) per μL of blood following ART release throughout the N-803 phase. Results are displayed for each vaccinated animal (blue) and control animal (red) individually.

**Supplemental Figure 4.**
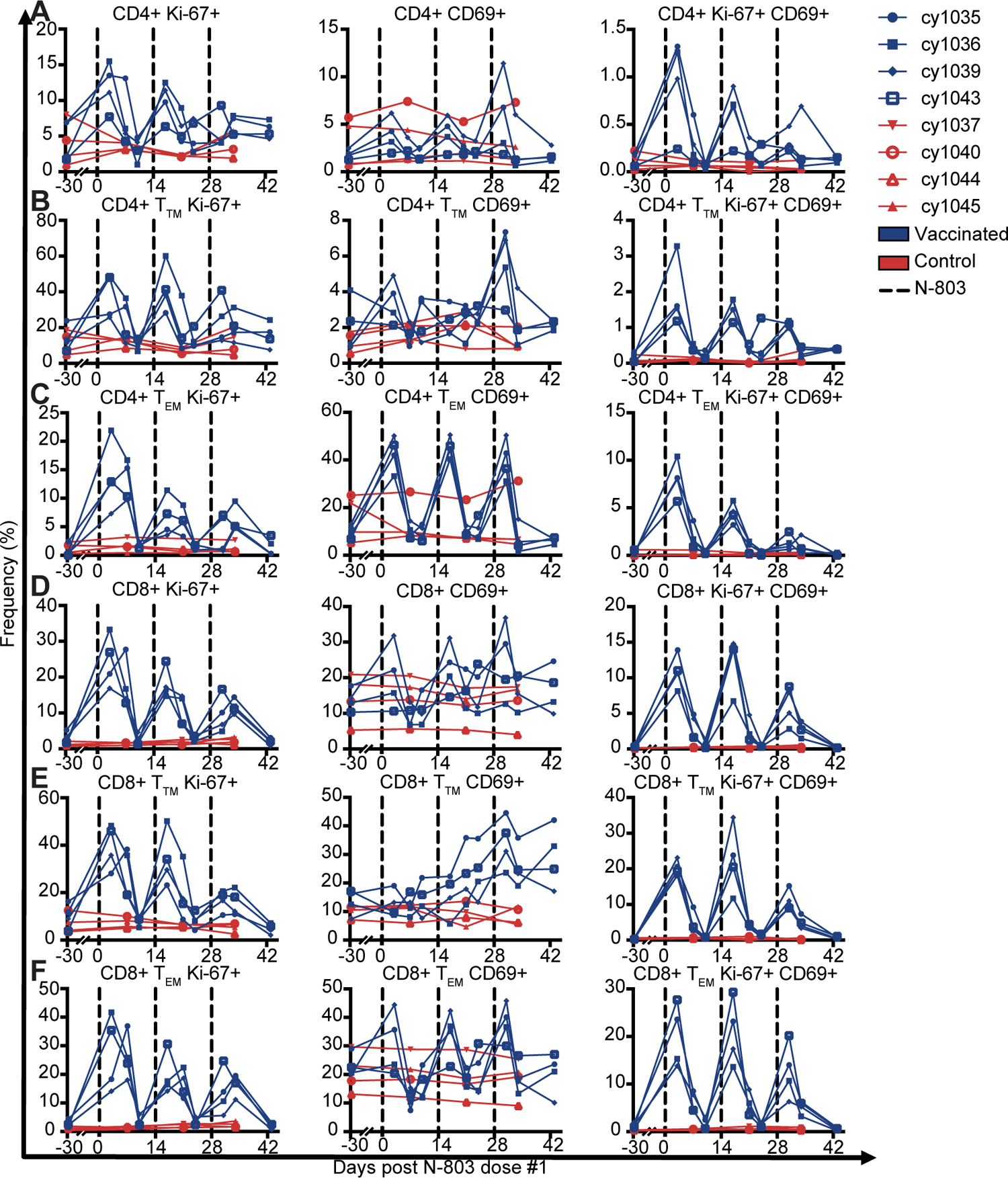
N-803-mediated expression of Ki-67 and/or CD69 by CD4+ and CD8+ bulk T cells, T_TM_, and T_EM_ in the PBMC. Frequency of **(A)** bulk CD4+ T cells, **(B)** CD4+ T_TM_, **(C)** CD4+ T_EM_, **(D)** bulk CD8+ T cells, **(E)** CD8+ T_TM_, and **(F)** CD8+ T_EM_ expressing the proliferation marker Ki-67 alone (left) the activation marker CD69 alone (center), or Ki-67 and CD69 together (right) throughout the N-803 phase. Results are displayed for each vaccinated (blue) and control (red) animal individually.

**Supplemental Figure 5.**
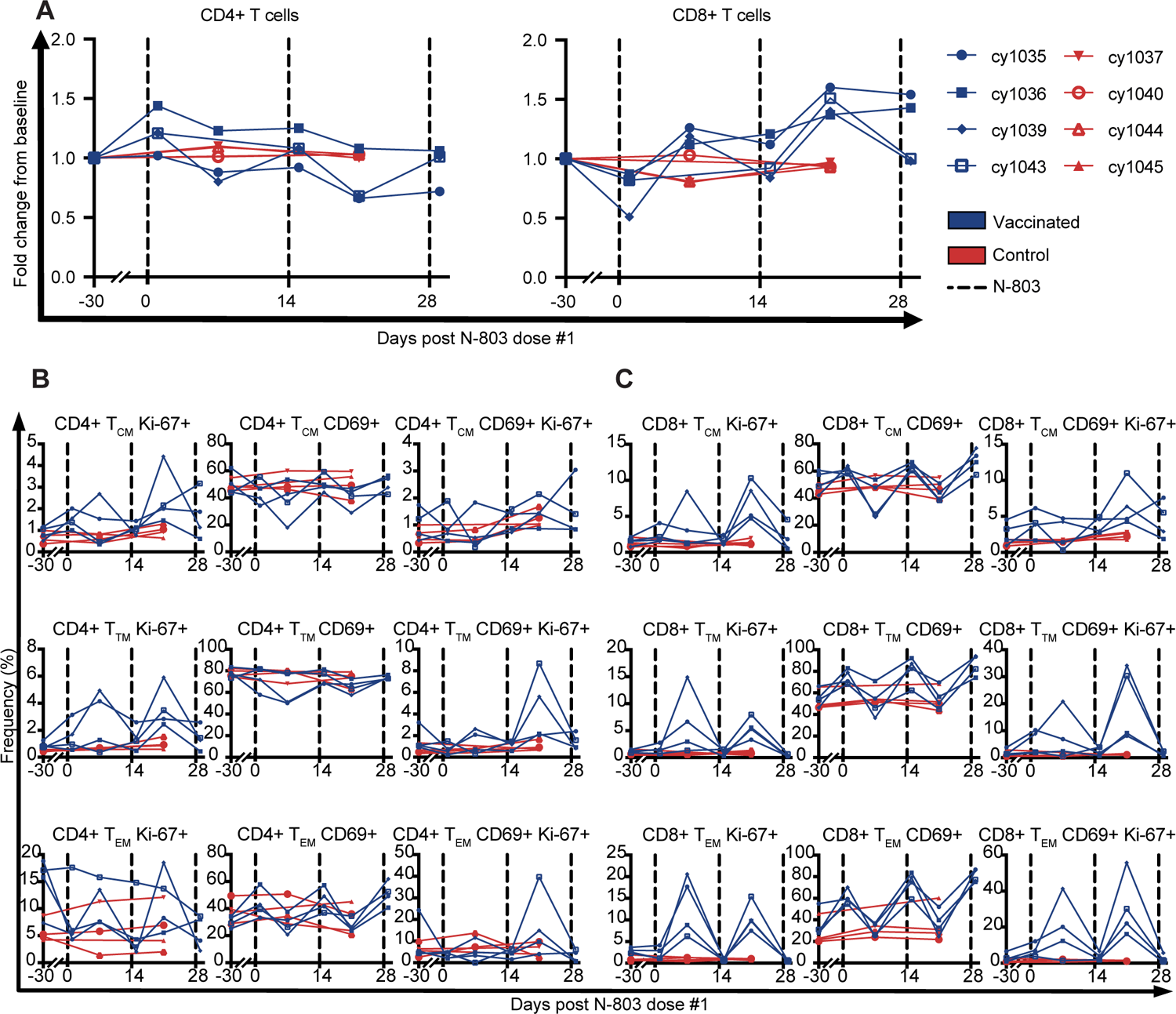
N-803-mediated fold changes of CD4+ and CD8+ T cells and expression of Ki-67 and/or CD69 by CD4+ and CD8+ T_CM_, T_TM_, and T_EM_ in the LN. (A) Bulk CD4+ T cells (left) and bulk CD8+ T cells (right). Results are displayed for each vaccinated (blue) and control (red) animal individually in fold change from baseline. **(B)** Frequency of CD4+ T_CM_ (top row), CD4+ T_TM_ (center row), and CD4+ T_EM_ (bottom row) expressing the proliferation marker Ki-67 alone (left), the activation marker CD69 alone (center), or Ki-67 and CD69 together (right) throughout the N-803 phase. **(C)** Frequency of CD8+ T_CM_ (top row), CD8+ T_TM_ (center row), and CD8+ T_EM_ (bottom row) expressing Ki-67 alone (left), CD69 alone (center), or Ki-67 and CD69 together (right) throughout the N-803 phase. Results are displayed for each vaccinated (blue) and control (red) animal individually.

